# A Chronnectogram-ICA Framework for Tracking Continuously Evolving and Overlapping Functional Brain States

**DOI:** 10.64898/2026.10.14.759150

**Authors:** N. Soleimani, S-L. Wiafe, A. Iraji, A. Preda, G. Pearlson, T. Adali, V. D. Calhoun

**Affiliations:** Department of Computer Science, Georgia State University, Atlanta, GA, 30303, USA; Department of Psychiatry and Human Behavior, University of California, Irvine, CA, 92697, USA; Departments of Psychiatry and Neuroscience, Yale University School of Medicine, New Haven, CT, 06511, USA; Department of Computer Science and Electrical Engineering, University of Maryland, Baltimore County, Baltimore, MD, 21250, USA; School of Electrical and Computer Engineering, Georgia Institute of Technology, Atlanta, GA 30332, USA

**Keywords:** Chronnectogram, Dynamic functional network connectivity (dFNC), Evolving states, Schizophrenia

## Abstract

The discrete-state model of brain dynamics has become the dominant paradigm in functional connectivity analysis, representing network activity as transitions among a fixed set of transiently recurring states while generally assuming that the states themselves do not evolve. We introduce a multilevel chronnectogram framework that relaxes this assumption and explicitly represents the higher-order temporal structure of brain dynamics, how distinct states couple and co-evolve over time, alongside the persistence of each state. The method integrates spatially guided independent component analysis (ICA) with subject-specific, state-guided dynamic ICA to capture evolving and overlapping connectivity patterns in a shared representational space. Validation on simulations with known ground truth shows recovery of continuous overlap and higher-order coupling structure invisible to discrete-state methods. Applied to resting-state functional magnetic resonance imaging from the FBIRN dataset (151 schizophrenia, 160 controls), the framework reveals that the same pair of brain states is shifted toward opposite extremes across patients, with strong synchrony in some and active anti-coupling in others, creating a redistribution of connectivity regimes invisible to conventional occupancy metrics, providing a substrate in which schizophrenia can appear both hyper- and hypo-volatile depending on the measure applied.

## 1 Introduction

The brain is a continuously evolving dynamical system, yet dominant paradigm for analyzing its time-varying functional organization commonly impose discrete connectivity states [1, 2]. Dynamic functional network connectivity (dFNC), a leading framework for studying time-resolved interactions among intrinsic connectivity networks (ICNs) [3], has become nearly synonymous with discretization; the moment-to-moment activity of the brain is reduced to a sequence of transitions among a small, fixed set of recurring connectivity states via clustering [3, 4], hidden Markov models (HMM) [5], or related state-assignment methods [6]. This abstraction has enabled tractable and reproducible characterization of dynamics across health and disease, but it reflects the constraints of available tools rather than an established property of neural dynamics. Whether neural dynamics truly occupy discrete states or evolve continuously remains unresolved, and committing to discreteness at the outset bounds what any subsequent analysis can recover.

By imposing hard state boundaries, conventional approaches discard the smooth transitions, partial overlaps, and short-lived configurations that may carry meaningful information [2, 7, 8], with consequences including reduced sensitivity to brief but informative fluctuations, a limited ability to model gradual transitions, and a vulnerability to oversimplification in high-dimensional feature spaces where coarse snapshots obscure subtle dynamics [7, 9]. A deeper limitation lies beneath these: discretization describes only how individual states are occupied over time and offers no representation of how distinct states relate to one another as they evolve, whether they co-activate, overlap, or move in opposition [8, 10].

Progress toward continuous representations has come largely from decomposition-based methods, which model functional imaging data as mixtures of latent sources rather than as assignments to discrete states [11,12]. Among these methods, independent component analysis (ICA) is a blind source separation technique that effectively identifies latent spatial networks from mixed fMRI signals, but its components are not consistently ordered or identified across subjects, making comparisons difficult [13–15]. Constrained ICA addresses this by incorporating spatial priors, such as templates from anatomical atlases or previously learned networks, enabling fully automated single-subject decompositions that retain correspondence across subjects [16–18]. Yet it has rarely been adapted to dynamic settings, and existing implementations are not designed to preserve fine-grained temporal fidelity while remaining aligned with group-level priors [17, 19]. No current approach recovers individualized, continuously evolving state trajectories while also representing how those states couple over time.

To close this gap, we introduce a chronnectogram-based framework that models the continuous evolution, overlap, and interaction of functional connectivity states within a unified representation. The framework unfolds as a multilevel decomposition. Spatially constrained ICA first extracts reproducible ICNs [19]; group-level connectivity ICA then identifies canonical connectivity states shared across the cohort, termed dynamic double functional independent primitives (ddFIPs) [20]; and a subject-specific dynamic connectivity-guided ICA recovers how each subject’s expression of those states evolves over time, building on reference-constrained ICA [17], yielding individualized evolving-state trajectories that remain aligned with group-level structure (Fig. 1). From these trajectories, we construct the chronnectogram, a state-by-state temporal coupling representation that encodes how distinct connectivity states co-evolve, succeed, and oppose one another over time, the higher-order structure that discrete-state methods leave unexpressed. Rather than assigning each window to a single state, the framework represents connectivity as a continuous trajectory in which states emerge, evolve, overlap, and interact.

**Figure 1:**
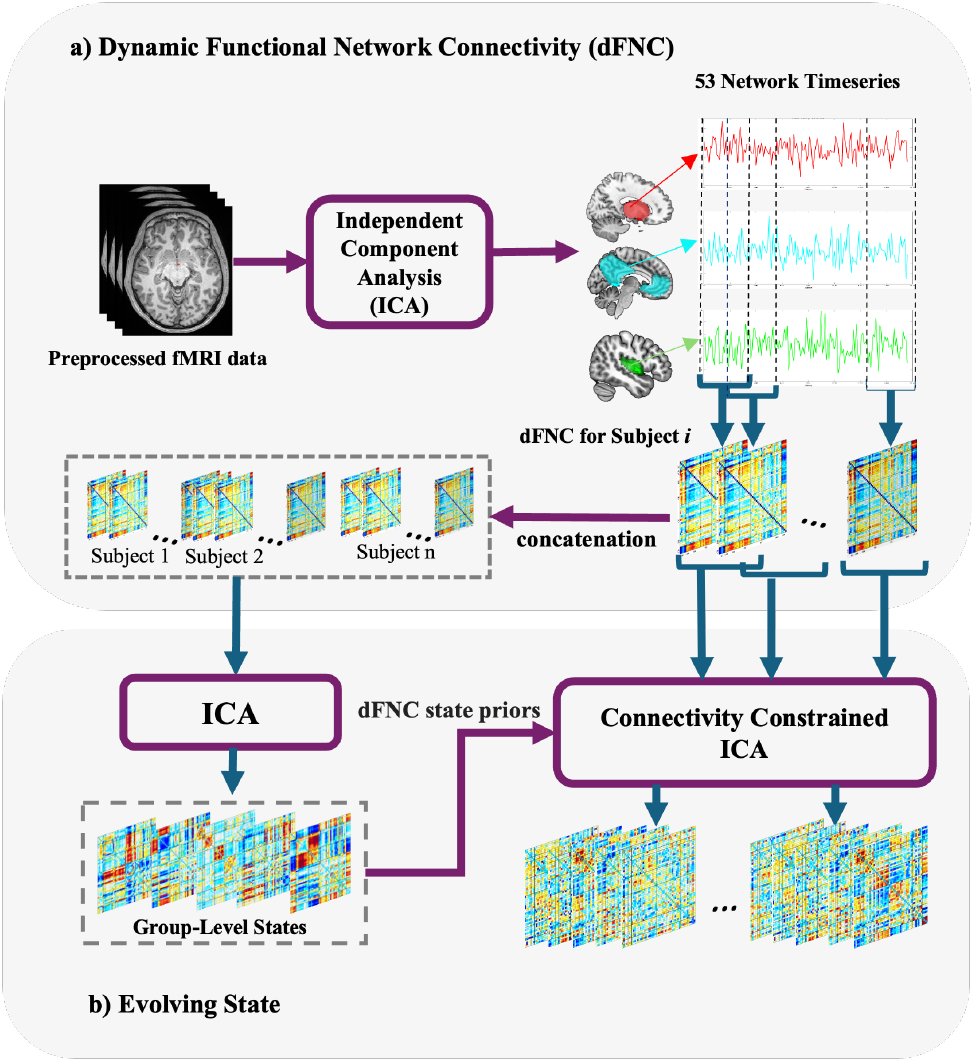
Schematic representation of the methodological pipeline for characterizing evolving states. (a) Extraction of ICN time series from preprocessed fMRI data, computation of dFNC using sliding-window correlations, and concatenation of subject-specific dFNC matrices. (b) Identification of group-level connectivity states via ICA, subsequently used as spatial priors in dynamically applying constrained ICA at the individual subject level, enabling detailed tracking of how connectivity states evolve temporally (evolving states).

We first validate the framework using simulations with known evolving states, overlap, and coupling in advance. It recovers continuous overlap and cross-state coupling, while remaining on par with discrete-state methods when the dynamics are genuinely discrete, so that its added expressiveness carries no penalty in the regime the discrete model was built for. It further detects group differences embedded purely in the distribution of state coupling, differences that leave first-order summaries unchanged and are therefore invisible to discrete-state analysis. In this sense, the framework does not replace the discrete description but generalizes it, with discreteness emerging as the limiting case of a continuous trajectory rather than an assumption imposed from the outset.

Applied to resting-state fMRI from the Functional Biomedical Informatics Research Network (FBIRN) cohort [21], it yields individualized trajectories of evolving connectivity and supports group-level inference on their coupling structure. Because the chronnectogram preserves the full distribution of state coupling rather than its average, it exposes effects that first-order summaries such as state occupancy and dwell times cannot reach. In schizophrenia (SCZ), it reveals state pairs driven toward opposite coupling extremes across patients, a redistribution that would be averaged away by mean-based metrics and that may reconcile reports of both increased and decreased dynamic variability. The contribution is therefore at once representational and inferential: a shift in the unit of analysis from the occupancy of individual states to the higher-order structure of how states couple as they evolve, and a distribution-aware basis for group inference that detects disease-related reorganization invisible to conventional dynamic connectivity metrics.

## 2 Methods

### 2.1 Data and Preprocessing

We evaluated our framework on a large multi-site dataset from the FBIRN study [21], including resting-state fMRI data collected from a total of 160 controls (45 female/115 male, average age: 37.04 ± 10.86 years, ranging from 19 to 59 years) and 151 individuals diagnosed with SCZ (36 female/115 male, average age: 38.77 ± 11.63 years, ranging from 18 to 62 years), meticulously matched in terms of age, sex distribution, and mean framewise displacement during scans (age: p = 0.18; sex: p = 0.39; mean framewise displacement: p = 0.97). Scans were collected at a repetition time (TR) of 2 seconds. All schizophrenic patients were in a clinically stable condition during the time of their scans. Standard preprocessing included slice-timing correction, motion realignment, normalization to MNI space (3*×*3*×*3 mm isotropic voxels), and spatial smoothing with a Gaussian kernel of a full width at half maximum (FWHM) of 6 mm. We applied NeuroMark spatially constrained ICA to extract 53 canonical ICNs, covering all major functional domains of the brain including subcortical (SC), auditory (AUD), sensorimotor (SM), visual (VS), cognitive control (CC), default mode (DM), and cerebellar (CB) networks for each subject, with detrending and despiking applied to enhance fMRI quality.

### 2.2 Extraction and Computation of dFNC

Following preprocessing, we extracted subject-specific time courses from *M* = 53 ICNs (Fig. 1). For subject *i*, let 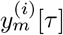 denote the time course of ICN *m* ∈ {1, …, *M*} at discrete time index *τ* ∈ {1, …, *T*}, where *T* is the total number of time points.

DFNC was estimated using a sliding-window Pearson correlation. Let *L* denote the window length (in number of time samples), for each window starting at index *t* (with 1≤*t*≤*T*−*L* + 1), we form the windowed correlation matrix

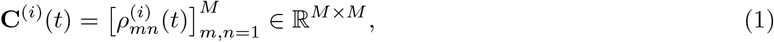

where 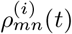 is the Pearson correlation between the windowed segments 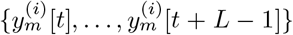 and 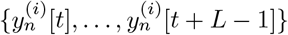

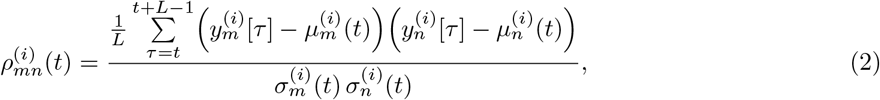

with the window-specific mean and standard deviation defined as

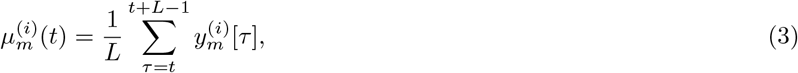

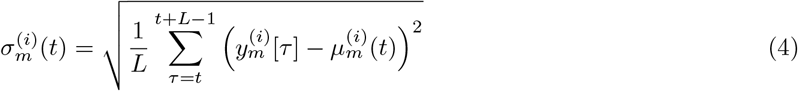

This formulation defines **C**(*t*) as a symmetric, positive semi-definite, covariance-normalized matrix that captures second-order dependencies between brain networks.

### 2.3 Group-Level Connectivity States

Let *N* denote the number of subjects and let *w* denote the number of sliding windows (dFNC samples) per subject. For subject *i* ∈ {1, …, *N* and window index *t* ∈ {1, …, *w*}, let **x**_*i*_(*t*) ∈ ℝ^*D*^ denote the vectorized dFNC feature vector obtained by stacking the (*D* = *M* (*M* −1)*/*2 = 1378) unique off-diagonal entries of the windowed correlation matrix **C**^(*i*)^(*t*).

We form the subject-level windowed dFNC matrix

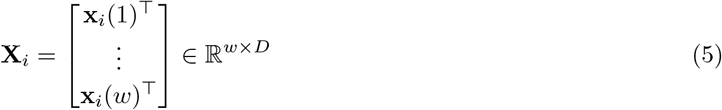

Concatenating all subjects along the window (row) dimension yields the group matrix

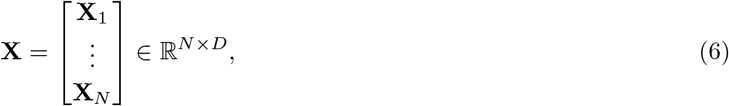

Group-level connectivity states were estimated by applying Infomax ICA (implemented in the GIFT toolbox) to **X** (after standard centering and whitening). In the Infomax formulation [22], an unmixing matrix **W** ∈ ℝ^*K×N*^ is estimated to maximize the joint entropy of the nonlinearly transformed outputs

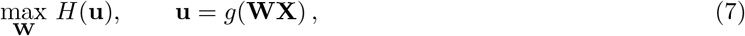

where *g*(*·*) is an element-wise sigmoid nonlinearity and

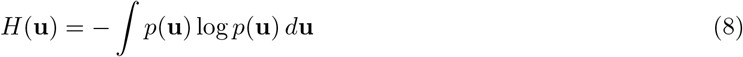

is the differential entropy of a continuous random vector **u**; *p*(**u**) denotes the probability density function of **u**.

Maximizing *H*(**u**) under the Infomax principle encourages each component of **u** to carry as much information as possible, while implicitly reducing mutual information between components. This yields statistically independent outputs corresponding to maximally independent patterns.

After convergence, the ICA model is written as

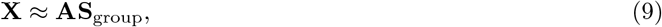

where **S**_group_ ∈ ℝ^*K×D*^ contains *K* group-level connectivity states, and **A** ∈ ℝ^*N ×K*^ contains the corresponding mixing coefficients across concatenated windows. The model order *K* was selected using the elbow criterion on the PCA eigenvalue spectrum, balancing model complexity and explanatory power; here we set *K* = 5.

### 2.4 Subject-Specific Dynamic Evolving States

To recover subject-specific states while preserving correspondence to group-level connectivity patterns, we applied multi-objective optimization ICA with reference (MOO-ICAR) [19] to each subject’s windowed dFNC matrix **X**_*i*_ ∈ ℝ^*w×D*^. Let 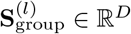 denote the *l*-th group-level state pattern, for *l* ∈ {1, …,}. For each subject *i* and state *l*, MOO-ICAR estimates an unmixing vector 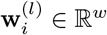 by solving

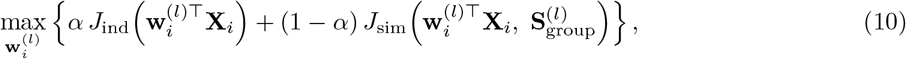

subject to 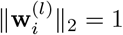. The resulting subject-specific state estimate is

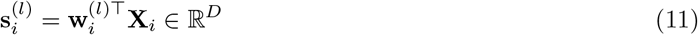

Here, *J*_ind_(*·*) promotes statistical independence (e.g., via negentropy/non-Gaussianity) and *J*_sim_(*·,·*) enforces similarity to the group level prior (e.g., correlation or mutual information). The parameter *α* ∈ (0, 1) controls the trade-off between independence and correspondence.

To obtain time-resolved evolving state trajectories, we apply the constrained decomposition across the sliding-window dFNC sequence using window length *L* = 40 samples and step size *s* = 2 samples, yielding *w* = 49 windows per subject. This configuration achieves a favorable trade-off between temporal resolution and stability of correlation estimates, with 90% overlap promoting smooth temporal tracking [3]. The resulting evolving representation for subject *i* and state *l* is denoted by

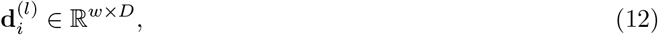

The resulting time courses and states reflect each subject’s expression of the canonical states evolves over time. Fig. 2 illustrates this evolution for a representative SCZ patient, showing activation of state 3 at selected time windows (10, 20, 30, and 40). Although all subjects share the same reference states, their temporal expression patterns are individualized. The supplementary Video presents the complete temporal evolution of all connectivity states for this exemplar patient.

**Figure 2:**
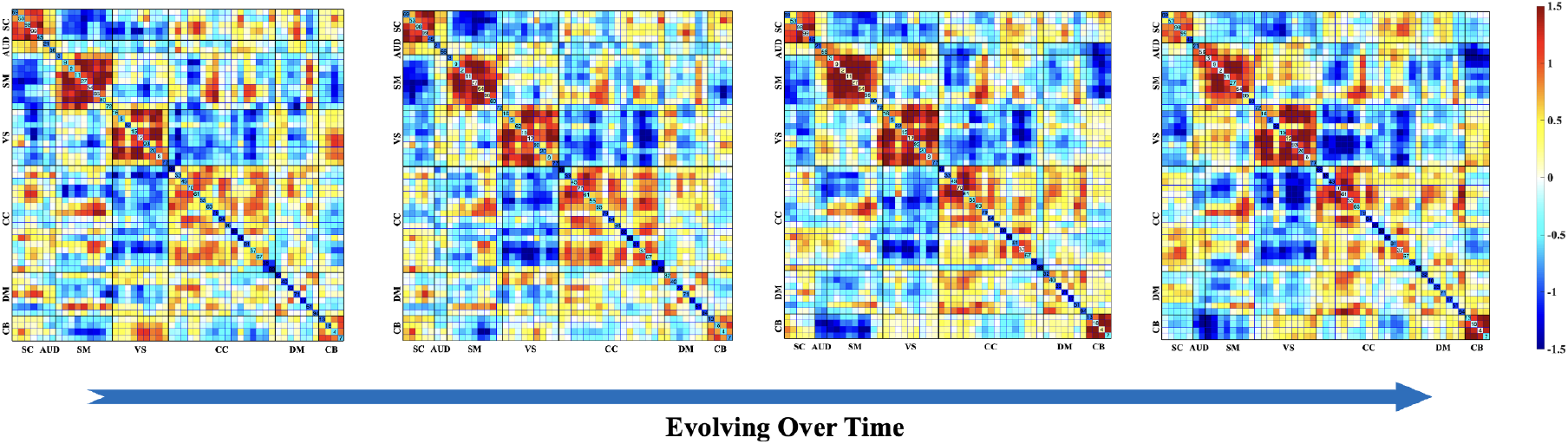
Visualization of temporal evolution of the third connectivity state across selected temporal windows (10, 20, 30 and 40) for an exemplar SCZ patient. The figure demonstrates how connectivity patterns dynamically evolve over time, highlighting transient changes and state transitions captured by the proposed framework. Supplementary Video provides an animation of the complete evolution of all connectivity states for this participant.

### 2.5 Inter-Subject Synchronization of Evolving States

To quantify synchrony of evolving state trajectories across subjects, we computed inter-subject similarity matrices for each state *l* ∈ {1, …, *K*}.

We define the mean connectivity pattern for subject *i* and state *l* across all windows as

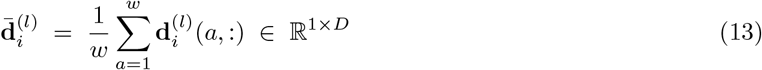

For subjects *i* and *j* and window indices *a, b* ∈ {1, …, *w*}, the inter-subject similarity at window-pair (*a, b*) is computed as the Pearson correlation across the *D* connectivity features:

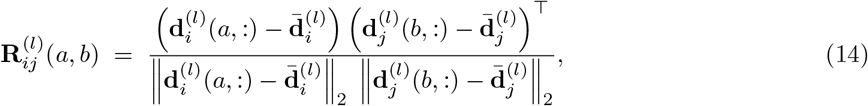

where ∥ · ∥_2_ denotes the Euclidean norm. The resulting matrix 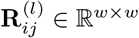 summarizes the correspondence between the temporal evolution of state-*l* connectivity patterns for the subject pair (*i, j*) across all window pairs.

We compute 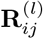 for within-group (HC–HC, SCZ–SCZ) and cross-group (HC–SCZ) subject pairs, and summarize each matrix using its maximum and minimum values to identify unusually high or low dynamic similarity pairs.

### 2.6 Chronnectogram Construction and Intra-Subject State Coupling

To characterize intra-subject interactions between dynamic connectivity states, we constructed chronnectograms that capture the temporal co-evolution of state-specific connectivity patterns.

To quantify temporal correspondence between two states *l*_*p*_ and *l*_*q*_ ∈ {1, …, *K*} and subject *i*, we computed the correlation between their evolving state trajectories, resulting in the chronnectogram:

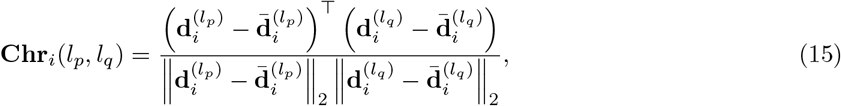

The resulting matrix **Chr**_*i*_(*l*_*p*_, *l*_*q*_) ∈ ℝ^*w×w*^ encodes correlations between the evolving connectivity profiles of the two states as shown in Fig. 4. Chronnectogram provides a structured representation of how different states interact and evolve over time by revealing synchronous, sequential, or antagonistic cross-state dynamics within an individual.

### 2.7 Chronnectogram Derived Histograms

Direct comparisons of chronnectograms across subjects are confounded by temporal misalignment in state transitions. To address this, we adopted a distribution-based approach by summarizing each chronnectogram as a two-dimensional histogram.

Formally, let 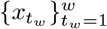 and 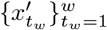 be the temporal trajectories of two connectivity states, the 2D histogram 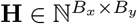 is defined as:

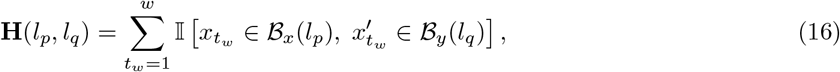

where ℬ_*x*_(*l*_*p*_) and ℬ_*y*_(*l*_*q*_) are the *l*_*p*_-th and *l*_*q*_-th bins along the *x*- and *y*-axes, respectively, and I[ ·] is the indicator function that equals 1 when the condition is true and 0 otherwise.

This histogram provides a compact, distribution-based representation of how frequently different coupling regimes occur for a given state pair, while reducing sensitivity to the exact timing of transitions.

For group analysis, two-sample *t*-tests were performed independently for each histogram bin to compare HC and SCZ (with false discovery rate correction for multiple comparisons). Within the SCZ cohort, associations between histogram features and clinical outcomes were assessed using general linear models with age and sex as covariates, including PANSS positive/negative and CMINDS composite cognition.

### 2.8 Simulation-Based Validation with Known Ground Truth

Because empirical fMRI provides no ground truth for assessing recovery, we conducted controlled Monte Carlo simulations in connectivity space, where evolving states, their overlap, and temporal coupling were specified a priori. This design isolates trajectory recovery from upstream preprocessing effects that could otherwise confound recovery failures. Synthetic windowed dFNC trajectories were generated with controllable state overlap, signal-to-noise ratio (SNR), and group-level differences, and were benchmarked against *k*-means and a Gaussian HMM.

We synthesized *K* = 5 neutral ground-truth states **B** ∈ ℝ^*K×D*^ as sparse (20% nonzero), FC-like patterns with bounded mutual similarity (maximum |*r*| = 0.2). Per-window state expression, **A**_*i*_ ∈ℝ^*w×K*^ (*w* = 49), followed two regimes: discrete (one-hot Markov chain, self-transition 0.9, zero overlap, the regime for which hard-label methods are designed), and continuous (a smoothly evolving simplex trajectory, the softmax of an Ornstein-Uhlenbeck process at time scale *τ* = 10, a temperature parameter sweeping overlap from zero to near-uniform). Windowed dFNC was generated as

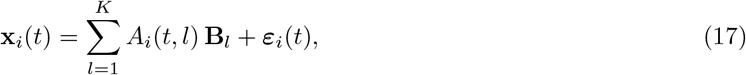

at a SNR of 2.

All three methods received the same group structure **B** as initialization so that representational capacity alone distinguished performance. The proposed estimator was initialized using its reference-similarity objective, and the *k*-means and Gaussian HMM baselines were initialized at **B** through their centroids and emission means, respectively. Any residual gap therefore reflects graded versus hard state membership, not privileged priors. All methods received identical data at the same model order, and recovered states were matched to ground truth via the Hungarian algorithm on absolute correlation.

The HMM used Gaussian emissions on a PCA-reduced representation of the windowed dFNC (*K* + 3 components for stable covariance estimation) with full Baum-Welch re-estimation. To confirm that its behavior reflects the model class rather than covariance handling, we swept the emission covariance structure across four configurations (spherical, diagonal, tied, and full) and separately varied posterior softness via temperature-scaled responsibilities to both the fitted emission means and the true vertices **B** (Fig. 3); results were consistent across all configurations.

**Figure 3:**
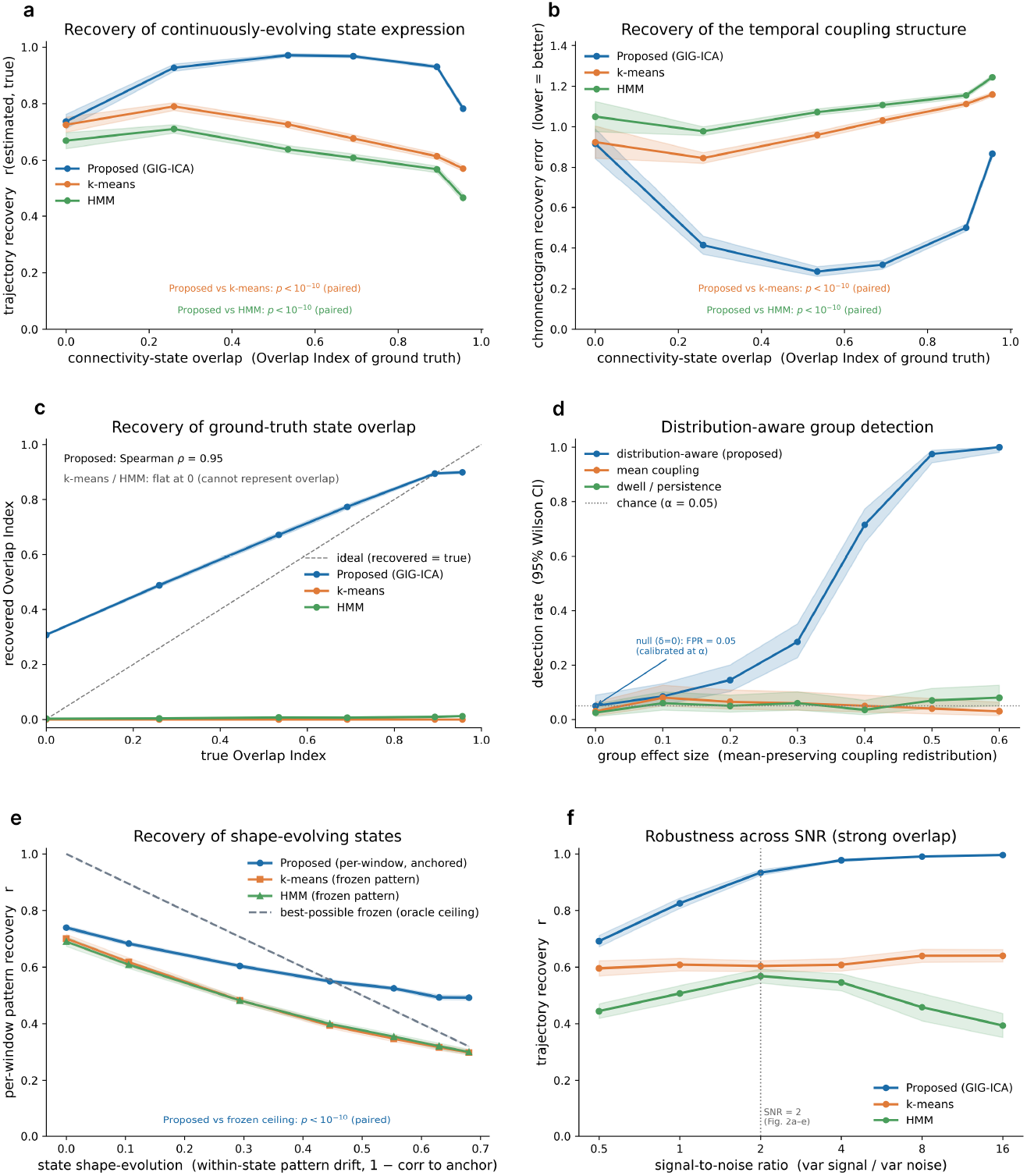
Chronnectogram-based evolving state recovery outperforms discrete assignment under continuous, overlapping brain dynamics. All methods are initialized at the true states so that any separation reflects representational capacity alone; lines are means over 200 simulated datasets, and bands are 95% confidence intervals. (a) Trajectory recovery as a function of ground-truth connectivity-state overlap: the methods are indistinguishable at zero overlap diverge with increasing overlap, and converge at near-uniform overlap. (b) Chronnectogram recovery error across the same overlap range. The proposed method achieves the lowest error at every level, with the largest margin at intermediate overlap. (c) Recovered versus true OI: the proposed method tracks ground-truth overlap monotonically (Spearman *ρ* = 0.95) while both baselines remain pinned at zero, structurally unable to represent simultaneous multi-state co-expression. (d) Detection rate of a planted mean-preserving group difference versus effect size: the distribution-aware test approaches certain detection while mean-coupling and dwell-time tests remain at chance, with false positives calibrated at the null (*FPR* = 0.05 at *δ* = 0). (e) Per-window pattern recovery under a shape-evolving generator: single-pattern methods are bounded by a decaying oracle ceiling (dashed) that the proposed estimator surpasses once drift is substantial. (f) Trajectory recovery across signal-to-noise ratios at strong overlap; the vertical dashed line marks SNR = 2, the operating point for panels (a–e). The proposed method remains stable across the full range while the HMM degrades as likelihoods sharpen toward harder state assignments. Paired comparisons in (a), (b), and (e) reach *p <* 10^−10^.

We quantified: (i) trajectory recovery, the mean absolute correlation between recovered and true per-window state expression; (ii) chronnectogram recovery error, the relative Frobenius error of the state-pair temporal-coupling tensor; (iii) the Overlap Index (OI), an entropy-based effective number of co-active states; and (iv) per-window pattern recovery, evaluated under a shape-evolving generator in which each state’s connectivity pattern drifts smoothly across windows. Group sensitivity was assessed by planting a mean-preserving bimodal redistribution of state-pair coupling and comparing a distribution-aware second-order statistic against mean-coupling and dwell-time tests, with the null (*δ* = 0) verifying false-positive calibration. Robustness was tested by sweeping SNR from 0.5 to 16 and repeating the overlap analysis under a bounded, heteroscedastic noise model (Gaussian perturbation in Fisher-z space mapped through tanh), which reproduces real sliding-window variance structure without additional model-order parameters. Each condition comprised 200 independent datasets (N = 100 per group for detection); we report 95% confidence intervals and paired Wilcoxon signed-rank tests, emphasizing effect sizes over p-values.

## 3 Results

### 3.1 Ground-Truth Recovery of Evolving States

Under matched priors, all methods performed similarly at zero overlap, where hard labels are correct (trajectory recovery r = 0.74, 0.73, and 0.67 for the proposed method, k-means, and the HMM respectively; Fig. 3 (a)). The proposed method’s advantage emerged and widened as states overlapped, reaching r = 0.78 (95% CI [0.77, 0.79]) at strong overlap versus 0.57 and 0.47 for the baselines, before converging at near-uniform overlap where all representations become equally challenged. Since all methods were warm-started at the true states, this gap is representational: one-hot loadings cannot encode graded membership.

The chronnectogram recovery error showed the same pattern (Fig. 3 (b)). Methods were comparable at zero overlap, whereas the proposed method achieved the lowest error across overlap levels, with the largest margin in the mid-overlap regime (*≈* 0.30 versus *≈* 1.0 for the baselines), again converging at near-uniform overlap. The recovered OI tracked ground truth monotonically for the proposed method (Spearman *ρ* = 0.95), while both baselines remained pinned at zero across the full range, being structurally unable to represent the simultaneous co-expression of multiple connectivity states (Fig. 3 (c)). The distribution-aware test detected the planted mean-preserving group difference with a detection rate approaching 1.0 across effect sizes, while the mean-coupling and dwell-time tests remained at the nominal level throughout, with false positives calibrated at the null (*FPR* = 0.05 at *δ* = 0; Fig. 3 (d)). This is an existence proof that chronnectogram-level group differences can escape first-order summaries entirely. Under a shape-evolving generator, any single-pattern-per-state method is bounded by a ceiling that decays as states drift. The proposed per-window estimator is not subject to this bound and surpassed the frozen-representation ceiling at strong drift (*r* = 0.49 versus 0.32 and 0.30 for the baselines; Fig. 3 (e)), confirming that per-window reference anchoring captures morphing state structure that frozen representations cannot reach. The HMM’s failure was one of vertex recovery rather than softness. Even under full posterior softening, recovery plateaued well below the proposed method and only resolved when the true mixing vertices were supplied directly. The same geometry explains why the HMM degraded as SNR increased: sharper likelihoods drive harder state assignments, collapsing the posteriors further (Fig. 3 (f)). The proposed method’s advantage persisted robustly across SNR from 0.5 to 16, with baseline performance bounded by representational capacity rather than noise at every level tested. The advantage persisted robustly across SNR from 0.5 to 16 (Fig. 3 (f)), with baseline performance bounded by representational capacity rather than noise at every SNR level tested.

### 3.2 Qualitative Chronnectogram Behavior

The chronnectogram of a representative healthy control (Fig. 4) shows that connectivity dynamics are clearly state-dependent: In this figure, each within-state panel (*l, l*) is a 49*×*49 autocorrelation-style similarity matrix across windows. Some states (e.g., states 3-5) exhibit strong temporal persistence with slow drift, while others (states 1-4) are more transient, and cross-state interactions range from strong co-evolution (state 3–5) to persistently weak or temporally localized coupling (e.g., state 2–3).

**Figure 4:**
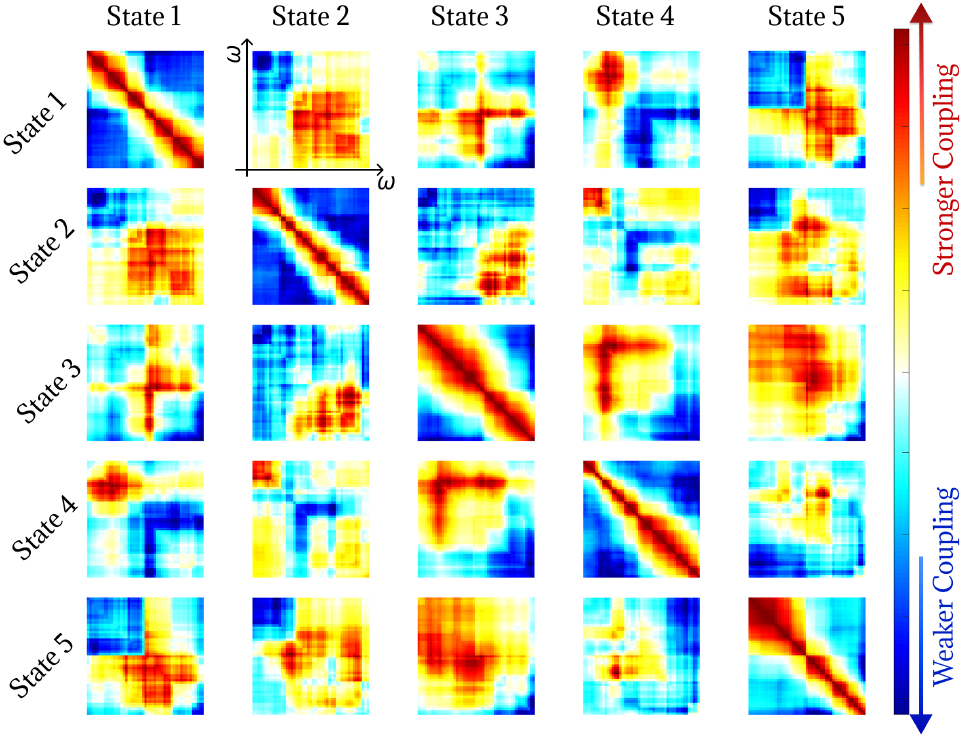
Chronnectogram of a representative HC individual. This matrix depicts dynamic temporal correlations between evolving states, highlighting how functional network configurations persist or transition. Warmer colors along the diagonal indicate sustained activation of the same state across consecutive time windows, whereas structured patterns off the diagonal reveal frequent transitions or co-activations between specific states.

### 3.3 Temporal Coupling Signatures Distinguish Schizophrenia and Controls

Using the chronnectogram-derived coupling histograms described elsewhere, we identified group differences in temporal coupling between SCZ and HC participants (Fig. 5), with warm and cool colors indicating SCZ- and HC-enriched coupling, respectively.

**Figure 5:**
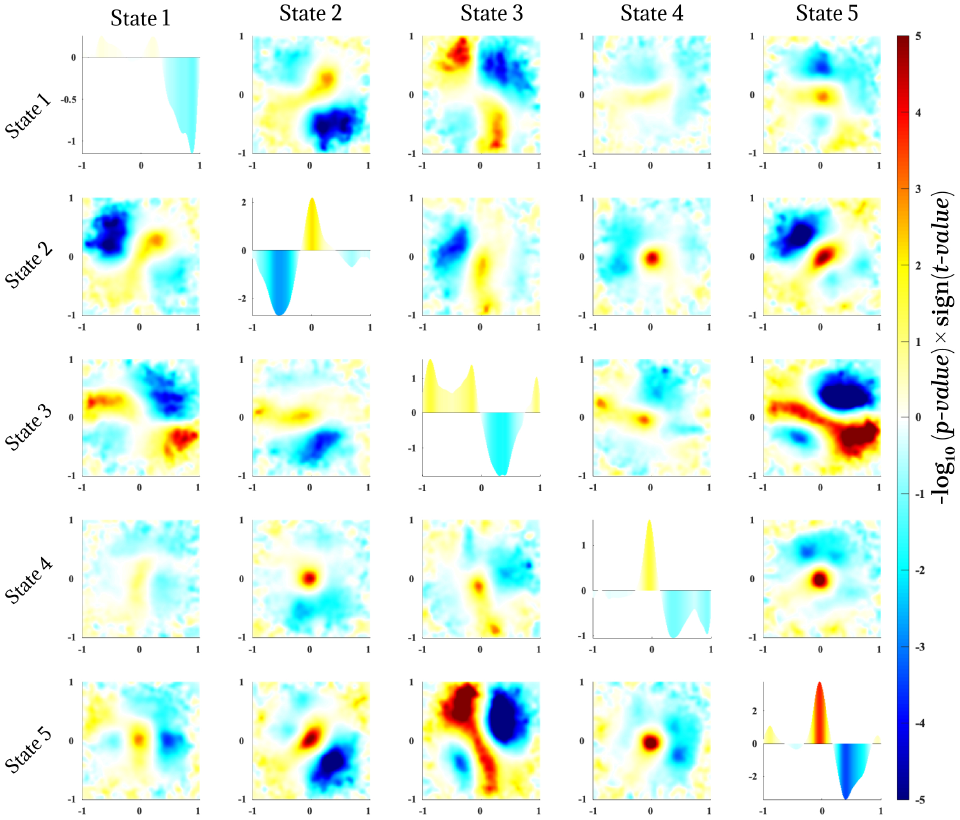
Group comparison of dynamic connectivity patterns between SCZ patients and healthy group, based on chronnectogram-derived coupling histograms. Each subplot represents a specific pair of connectivity states, with color indicating the *t*-statistic for the HC vs. SCZ difference at each level of coupling (self- or cross-correlation). Warm colors (positive *t*) denote stronger or more frequent coupling in SCZs relative to HCs, while cool colors (negative *t*) denote stronger coupling in HC group. Notable differences are seen both on the diagonal (within-state temporal stability) and off-diagonals (cross-state interactions), highlighting specific states and state pairs where dynamic connectivity diverges between groups.

Within-state effects were most prominent for state two, which showed a broad HC-enriched portion of the coupling distribution alongside a more localized SCZ-enriched region, suggesting reduced temporal persistence and greater volatility in SCZ. Cross-state comparisons revealed localized, state-pair-specific differences rather than uniform shifts: the state 1-state 2 and state 1-state 3 interactions both showed opposing-regime patterns at high–low coupling combinations, reflecting a negative relationship between the respective state trajectories.

The most striking effect, and one inaccessible to any first-order analysis, appeared in the state 3-state 5 interaction: the significance map was bimodal, with the SCZ group enriched at both high and low coupling extremes simultaneously. Some patients showed strong co-activation between these states, others a pronounced inverse relationship in which one state’s engagement excluded the other. Because these opposing shifts occupy opposite ends of the coupling distribution, they cancel under mean-based comparison and are invisible to occupancy or dwell-time summaries-yet the chronnectogram resolves them directly. This is the empirical signature the framework is designed to expose: a redistribution of dynamic coupling that conventional state metrics cannot represent.

### 3.4 Dynamic Expression Profiles at State Minima and Maxima

To test whether SCZ and HC participants differ in how dynamic connectivity states attenuate and peak over time, we analyzed inter-subject synchronization extrema, defined as the window indices corresponding to minimum and maximum coupling between subjects. For each state, extrema samples were clustered using *k*-means (*k* = 3) and the clustering was performed separately for minima and maxima. Group-level differences were then assessed by comparing the distributions of cluster membership across SCZ and HC cohorts (Fig. 6).

**Figure 6:**
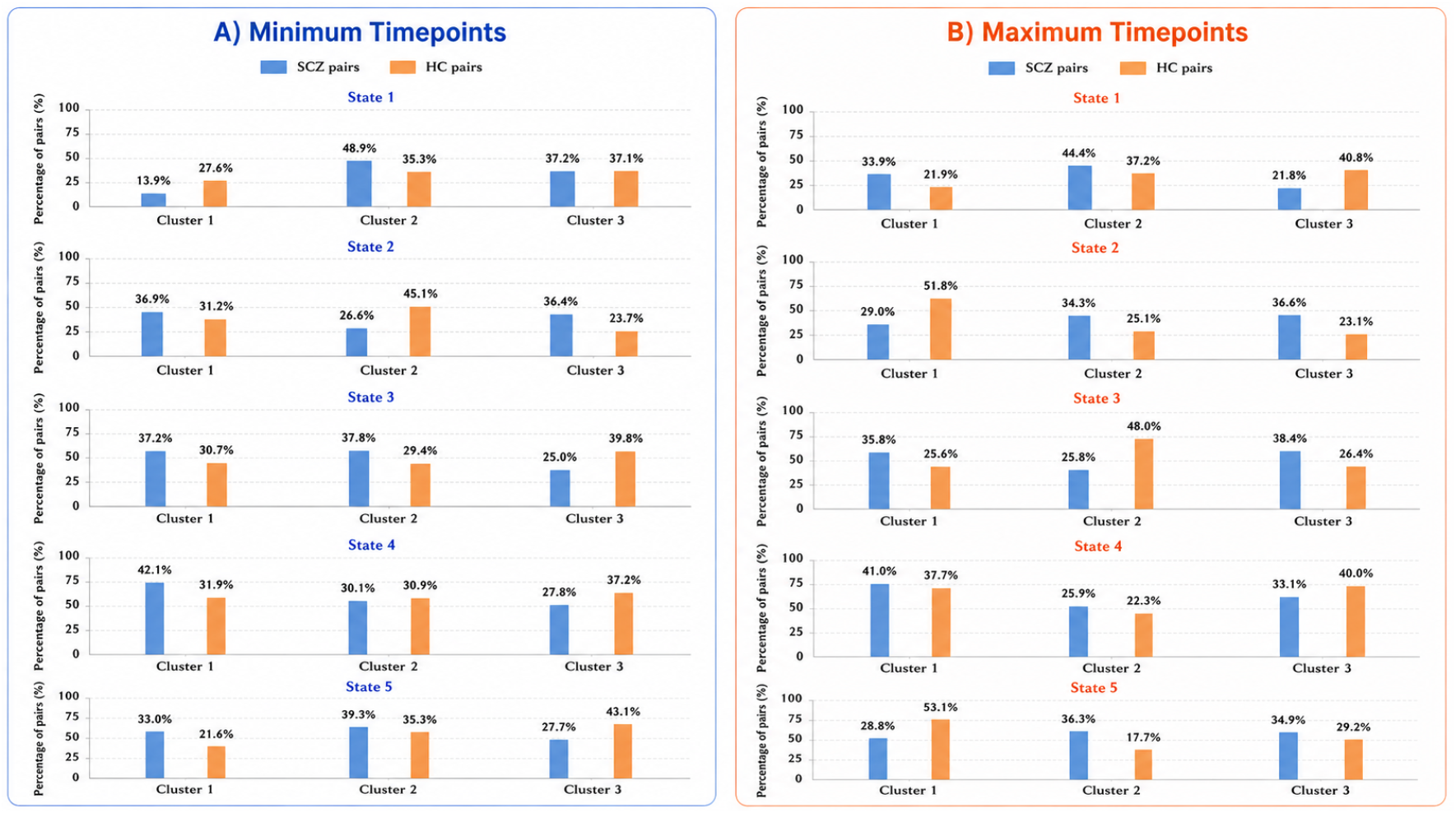
Cluster distributions of SCZ and HC subject pairs across dynamic states from *k*-means clustering of minimum and maximum time points (*k* = 3). Bar heights denote the percentage of pairs assigned to each cluster. SCZ pairs were more frequently assigned to distinct or less stable temporal configurations, whereas HC pairs more often showed integrated or synchronized dynamics.

Clear group-dependent structure was observed at both ends of the coupling spectrum. At minimum coupling, SCZ participants were systematically overrepresented in clusters corresponding to atypical low-coupling profiles. For instance, in state 1, approximately 48.9% of SCZ minima were assigned to a single cluster (Cluster 2) that was infrequently occupied by HCs, indicating a pronounced shift in the state-disengagement regime. A similar pattern was evident in state 2, where HC minima were concentrated within a dominant cluster, whereas SCZ minima were distributed across multiple clusters, consistent with increased heterogeneity in low-coupling configurations.

Analyses of maximum coupling further demonstrated dissociable group structure. In states exhibiting broader integrative connectivity, most notably state 5, HC participants preferentially occupied clusters characterized by relatively consistent high-coupling profiles, whereas SCZ participants were more frequently assigned to clusters reflecting less coordinated or atypical peak configurations. This distributional shift suggests that group differences extend beyond state weakening to include altered organization of peak state expression.

Collectively, the extrema-based clustering results indicate that SCZ is associated with systematic alterations in both state disengagement (minima) and state engagement (maxima). Relative to controls, the SCZ cohort exhibits a more heterogeneous distribution of extrema profiles, consistent with reduced temporal regularity in transitions into and out of dominant connectivity configurations.

### 3.5 Associations Between State Dynamics and Clinical Outcomes

We next evaluated whether inter-individual variability in chronnectogram-derived coupling features within the SCZ cohort covaries with symptom burden and cognitive performance. For each state pair, we fit general linear models (GLMs) with age and sex as covariates and tested the association between each coupling feature and three clinical measures: positive and negative symptom scores (PANSS) and computerized multiphasic interactive neurocognitive system (CMINDS). The resulting maps (Fig. 7 (a), Fig. 7 (b) and Fig. 7 (c)) display the signed strength of association at each coupling bin, where warm and cool values denote increased coupling expression associated with higher and lower clinical scores, respectively.

**Figure 7:**
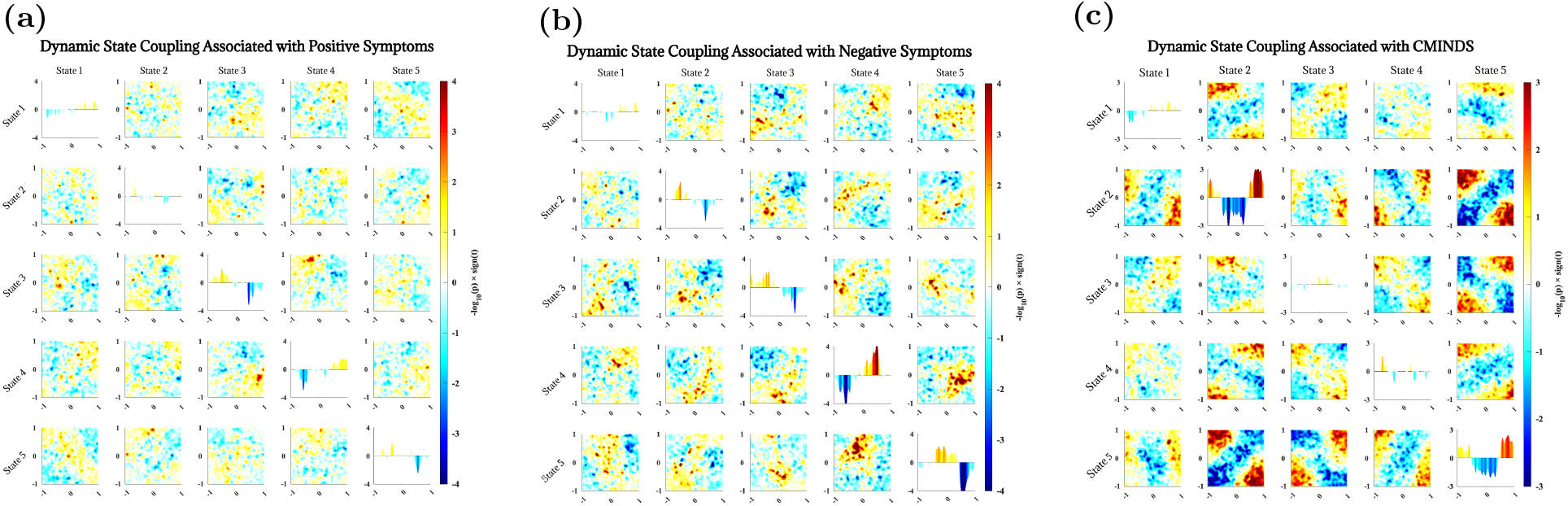
GLM associations between state-coupling features and symptom severity/clinical outcomes in SCZ. Colors indicate signed association strength, −log_10_(*p*) *·*sign(*t*); warm and cool values denote stronger positive and negative associations, respectively. (a) Positive symptoms: state 3-state 4 synchrony (black box) was associated with greater symptom severity, whereas anti-coupling was associated with lower severity. (b)Negative symptoms: greater within-state persistence in states 3–5, imbalanced state 4-state 5 coupling, and anti-correlated state 3-state 4 coupling were associated with greater severity. (c) CMINDS composite scores: cross-state interactions, particularly state 2-state 5 and state 4-state 5, were predominantly positively associated with cognitive performance, whereas few within-state effects were observed, indicating that cognition is more strongly related to inter-state coordination than state persistence.

Positive symptoms were associated with bidirectional coupling patterns (Fig. 7(a)). Greater synchrony between states 3 and 4 was associated with more severe positive symptoms, whereas anti-coupling between the same states was associated with lower symptom scores. Similar trends for state 1-state 3 and state 2-state 3 suggest that altered interactions across functional domains, including abnormal DM-CC co-activation, may contribute to positive-symptom expression.

Negative symptoms showed a distinct profile (Fig. 7(b)). Greater persistence within states 3, 4, and 5, imbalanced state 4-state 5 interactions, and anti-correlated state 3-state 4 coupling were associated with increased negative-symptom severity.

Cognitive performance was associated primarily with coordinated cross-state dynamics rather than within-state persistence (Fig. 7 (c)). Stronger co-activation of state 5 with state 2 (e.g., VS and SM) and State 4 (e.g., CC) predicted higher CMINDS scores, consistent with the importance of flexible large-scale network integration for preserved cognition. Conversely, coupling regimes in which state 2 activation accompanied state 5 suppression were associated with poorer cognitive performance, suggesting that excessive segregation between synergistic states may impair task-relevant integration.

Taken together, the GLM analyses demonstrate that symptom dimensions and cognition are associated with distinct regions of the state-coupling space. In technical terms, the chronnectogram-derived representation captures clinically relevant variability through regime-specific, bin-localized changes in the empirical coupling distribution, rather than through global shifts in mean coupling alone.

## 4 Discussion

In this study, we introduced and evaluated the *chronnectogram* as a representation of the higher-order temporal organization of evolving connectivity states. Rather than reducing dFNC to discrete state labels and first-order summary measures, the proposed framework characterizes the full temporal coupling structure between evolving states for each subject and compares the resulting empirical coupling distributions across groups. Applied to the FBIRN resting-state fMRI dataset, we revealed that SCZ is associated with selective alterations in within-state persistence, cross-state coupling, and extrema organization-expressed not as uniform shifts in mean coupling, but as state-specific redistributions across distinct regions of the coupling space. The most prominent group effects involved altered self-similarity of state 2, redistribution of cross-state coupling for pairs such as state 1-state 3 and state 3-state 5, increased heterogeneity in minima and maxima of state expression, and regime-specific associations with symptom and cognitive measures. Taken together, these findings indicate that the temporal organization of evolving network states is disrupted in SCZ at multiple levels of description, extending prior observations of abnormal state occupancy and transition structure in SCZ [10, 23–25].

### 4.1 Neural Coordination and Dynamic State Dysregulation in Schizophrenia

A central implication of findings is that SCZ-related dysconnectivity reflects an alteration in the occupancy of specific temporal coupling regimes rather than a simple change in overall connectivity strength. The chronnectogram-derived histograms showed that group differences were localized to specific state pairs and specific regions of the joint coupling space, indicating that disease effects reshape the empirical distribution of coupling values rather than merely shifting its mean. The altered diagonal structure, most evident for state 2, suggests reduced persistence of a subset of evolving connectivity configurations, while off-diagonal results show reorganized cross-state coordination, with some pairs exhibiting reduced positive coupling and others enhanced antagonism. The bimodal redistribution observed for state 3–state 5 is particularly informative, reflecting the emergence of distinct coupling regimes rather than a homogeneous group-level displacement. The present findings support the view that SCZ perturbs the balance among integrated, weakly coupled, and antagonistic network configurations over time, broadly consistent with prior dFNC studies reporting altered occupancy of synchronized and dysconnected states in SCZ [10, 23–26].

More broadly, these results are consistent with the interpretation of SCZ as a disorder of temporally dysregulated large-scale neural coordination. Healthy brain dynamics are thought to involve recurrent movement between relatively integrated and relatively segregated network configurations [2, 27], with transient high-synchrony regimes supporting large-scale coordination [28] and differentiated regimes supporting functional specialization. The redistribution of coupling regimes observed here suggests this balance is altered in SCZ, consistent with impaired large-scale integration [29, 30] and with recent accounts emphasizing abnormal coordination between integrated and segregated modes [31, 32]. This dysregulation was not uniform across symptom dimensions: positive symptoms were linked to bidirectional coupling regimes, negative symptoms to broader within-state persistence and selected cross-state interactions, and cognition primarily to coordinated cross-state coupling involving state 5. Clinical variance is therefore not encoded by a single dysconnectivity index, but by distinct regions of the state-coupling space.

### 4.2 Methodological Innovations over Traditional dFNC Approaches

The main methodological contribution is the integration of four complementary steps into a unified framework: (1) subject-level spatially guided ICA for reproducible ICN extraction, (2) group-level connectivity ICA for canonical state discovery, (3) subject-specific dynamic ICA for time-resolved individualized state estimation, and (4) chronnectogram-based analysis of higher-order temporal coupling between evolving states. This design addresses two limitations of conventional dFNC: hard assignment of windows to discrete states obscures continuously evolving or overlapping configurations [33], whereas our evolving-state representation allows expression to vary over time while remaining aligned with group-level priors [17, 19]. Further, classical summaries (occupancy, dwell time, switch counts) reduce dynamics to first-order statistics, while the chronnectogram preserves the full *w × w* within- and cross-state coupling structure, with the histogram summarizing this as an empirical distribution over coupling regimes.

This distinction is not merely representational; it directly affects what types of abnormalities can be detected. Many group differences identified here were bin-localized and, in some cases, reversed sign within the same state pair, indicating distributional rather than global effects-precisely the regime our simulations identified as invisible to first-order metrics but resolvable by the distribution-aware chronnectogram. A further technical advantage is that the histogram-based summary reduces sensitivity to exact temporal alignment across subjects while preserving information about over- or under-represented coupling regimes, making the method well suited to settings where disease-related effects may be expressed through changes in the distribution of state interactions rather than consistent absolute-time alignment. More generally, the framework bridges state-based dFNC analysis and trajectory-based descriptions of dynamic brain organization, an emphasis increasingly present in recent chronnectomic work [10, 33, 34].

### 4.3 Comparison with Previous Dynamic Connectivity Studies

The present results both corroborate and refine earlier dFNC evidence that SCZ-related abnormalities are state dependent and concentrated in specific connectivity configurations, including synchronized, weakly coupled, and antagonistic regimes [11, 23, 24, 35]. The chronnectogram framework extends this literature by localizing group effects within the joint coupling distribution of individual state pairs, rather than limiting inference to occupancy of discrete states.

This distributional perspective may also reconcile reports of both hyper- and hypo-volatility in SCZ [36,37]. Different conventional metrics may emphasize persistence, switching, or variability within the same underlying redistribution of coupling regimes. Our findings further indicate that SCZ is not defined by a single aberrant state, but by altered temporal coordination among multiple evolving states, including simultaneous increases and decreases in distinct coupling regimes within a state pair. By capturing these higher-order interactions, the chronnectogram bridges conventional state-based dFNC with trajectory-based models of network dynamics [33, 34].

### 4.4 Limitations and Future Direction

Several limitations merit consideration. The sliding-window correlation approach involves an inherent bias– variance tradeoff, and window parameters can influence the resulting state and chronnectogram structure [2,33]. Reliance on group-derived priors improves cross-subject correspondence but may bias estimates toward template-conforming patterns [17, 19]. Additionally, the histogram representation trades exact temporal sequencing for robustness to misalignment, making the framework well suited to characterizing coupling distributions but less informative about detailed state-sequence grammar. Finally, the FBIRN cohort’s clinical heterogeneity (medication status, illness duration) may contribute to observed patterns, so findings should be validated across independent cohorts, including first-episode and unmedicated samples [36].

In summary, by modeling chronnectogram-derived coupling distributions, the proposed framework captures higher-order network reorganization beyond discrete-state occupancy metrics and provides a basis for more informative dynamic biomarkers of psychiatric illness [10, 23, 33].

